# Analysis of the Human Connectome Data Supports the Notion of A “Common Model of Cognition” for Human and Human-Like Intelligence Across Domains

**DOI:** 10.1101/703777

**Authors:** Andrea Stocco, Catherine Sibert, Zoe Steine-Hanson, Natalie Koh, John E. Laird, Christian J. Lebiere, Paul Rosenbloom

## Abstract

The Common Model of Cognition (CMC) is a recently proposed, consensus architecture intended to capture decades of progress in cognitive science on modeling human and human-like intelligence. Because of the broad agreement around it and preliminary mappings of its components to specific brain areas, we hypothesized that the CMC could be a candidate model of the large-scale functional architecture of the human brain. To test this hypothesis, we analyzed functional MRI data from 200 participants and seven different tasks that cover a broad range of cognitive domains. The CMC components were identified with functionally homologous brain regions through canonical fMRI analysis, and their communication pathways were translated into predicted patterns of effective connectivity between regions. The resulting dynamic linear model was implemented and fitted using Dynamic Causal Modeling, and compared against six alternative brain architectures that had been previously proposed in the field of neuroscience (three hierarchical architectures and three hub-and-spoke architectures) using a Bayesian approach. The results show that, in all cases, the CMC vastly outperforms all other architectures, both within each domain and across all tasks. These findings suggest that a common set of architectural principles that could be used for artificial intelligence also underpins human brain function across multiple cognitive domains.

## Introduction

The fundamental organizational principle of a complex system is often referred to as its “architecture,” and represents an important conceptual tool to make sense of the relationship between a system’s function and structure. For instance, the von Neumann architecture describes the organizing principle of modern digital computers; it can be used both to describe a computer at a functional level of abstraction (ignoring the specific wiring of its motherboard and mapping it onto a theory of computation) and, conversely, to conduct diagnostics on an exceedingly complicated piece of hardware (properly identifying the components and pathways on a motherboard and the function of their wiring).

The stunning complexity of the human brain has inspired a search for a similar “brain architecture” that, akin to von Neumann’s, could relate its components to its functional properties. Succeeding in this quest would lead to a more fundamental understanding of brain function and dysfunction and, possibly, to new principles that could further the development of artificial intelligence (Hassabis et al., 2017).

Most attempts in this direction have been “bottom-up,” that is, driven by the application of dimensionality-reduction and machine-learning methods to large amounts of connectivity data, with the goal of identifying clusters of functionally connected areas (Cole et al., 2013; Gorgolewski et al., 2014; Huntenburg et al., 2018). Although these models can be used to predict task-related activity, they rely on large-scale connectivity and are fundamentally agnostic (or, at best, make a task-specific guess) as to the function of each network node. The results of such approaches are also dependent on the type of data and the methods applied. For instance, one researcher might focus on purely functional measures, such as task-based fMRI and the co-occurrence of activity across brain regions and domains; a second researcher, instead, might focus on spontaneous, resting-state activity and slow frequency time series correlations.

As recently pointed out (Jonas & Kording, 2017), none of these methods is guaranteed to converge and provide a functional explanation *from* the data. However, the same methods can be successfully used to test two or more models *against* the data via a “top-down” approach (Jonas & Kording, 2017). That is, given a candidate functional model of the brain, traditional connectivity methods can provide reliable answers as to its degree of fidelity to the empirical data and its performance compared to other models. A top-down approach, however, critically depends on having a likely and theoretically-motivated functional proposal for a brain architecture.

### The Common Model of Cognition

A promising candidate proposal is the Common Model of Cognition (CMC) (Laird et al., 2017). As the name implies, the CMC is a common set of organizing principles that summarize the similarities of multiple cognitive architectures that were developed over the course of five decades in the fields of cognitive psychology, artificial intelligence, and robotics. It is an architecture for *general* intelligence, in the sense that agents based on its principles should be capable of exhibiting rational and adaptive behavior across domains, rather than optimal behavior in a narrow domain.^1^ Because of its generality and consensus, it has been used as a guideline for designing cognitive agents (Mohan, n.d.). According to the CMC, agents exhibiting human-like intelligence share five functional components: a feature-based, declarative long-term memory, a buffer-based working memory, a system for the pattern-directed invocation of actions represented in procedural memory, and dedicated perception and action systems.

Working memory acts as the hub through which all of the other components communicate, with additional connections between perception and action (Figure 1A). The CMC also includes additional constraints on the mechanisms and representations that characterize each component’s functional properties.

**Figure 1.**
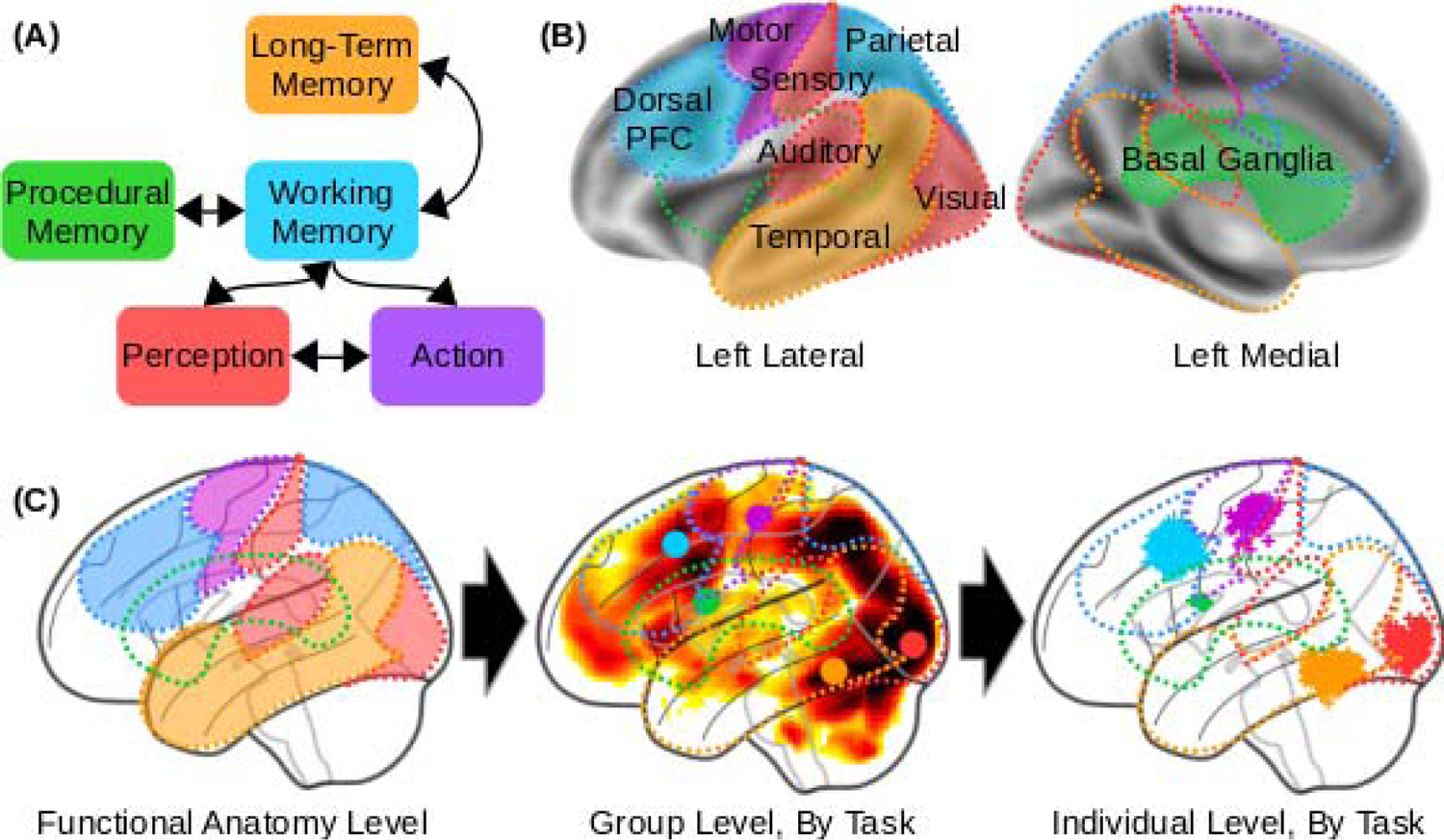
(A) Architecture of the Common Model of Cognition, as described by (Laird et al., 2017). (B) Theoretical mapping between CMC components and homologous cortical and subcortical regions, as used in this study’s pipeline to identify the equivalent Regions of Interest (ROIs). (C) Progressive approximation of the ROIs, from high-level functional mappings (left) to task-level group results (middle, with group-level centroid coordinated marked by a color circle) to the individual functional centroids of the regions in our sample (right; each individual centroid represented by a “+” marker; note that hundreds of markers are overlapping in each region). Group-level and individual-level data come from the Relational Reasoning task.

The CMC’s components and assumptions distill lessons learned over the last fifty years in the development of computational cognitive models and artificial agents with general human-like abilities. Surprisingly, these lessons seem to cut across the specific domains of application. For instance, the cognitive architecture Soar (Laird, 2012) is predominantly used in designing autonomous artificial agents and robots, while the cognitive architecture ACT-R (Anderson, 2007) is predominantly used to simulate psychological experiments and predict human behavior (Kotseruba & Tsotsos, 2020); yet, they separately converged on many of the CMC assumptions (Laird, Lebiere, & Rosenbloom, 2017). Similarly, the SPAUN large-scale brain model (Eliasmith et al., 2012) and the Leabra neural architecture (O’Reilly et al., 2016) are independently designed to simulate brain function through artificial neurons; despite making different assumptions in terms of neural coding, representation, and learning algorithms, they agree on the use of high-level modules (including ones for working memory, procedural memory, and long-term memory) that are similar to the CMC. Even recent AIs that are made possible by advances in artificial neural networks employ, at some level, the same components. DeepMind’s AlphaGo, for example, includes a Monte-Carlo search tree component for look-ahead search and planning (working memory) and a policy network (procedural memory), in addition to dedicated systems for perception and action (Silver et al., 2016). Similarly, the Differentiable Neural Computer (Graves et al., 2016) uses supervised methods to learn optimal policies (procedural memory) to access an external memory (symbolic long-term memory).

Because the CMC reflects the general organization of systems explicitly designed to achieve human-like flexibility and intelligence, the CMC should also apply to the human brain. Therefore, it provides an ideal candidate for a top-down examination of possible brain architecture.

### Assessing the CMC as a Brain Architecture

Assuming that the CMC is a valid candidate, how can its viability as a model of the human brain architecture be assessed? Operationally, a candidate model should successfully satisfy two criteria. The first is the *generality* criterion: the same cognitive architecture should account for brain activity data across a wide spectrum of domains and tasks. The second is the *comparative superiority* criterion: an ideal architecture should provide a superior fit to experimental brain data compared to competing architectures of similar complexity and generality.

To test the CMC against these two criteria, we conducted a comprehensive analysis of task-related neuroimaging data from 200 young adult participants in the Human Connectome Project (HCP), the largest existing repository of high-quality human neuroimaging data.

Although the HCP project contains both fMRI and MEG data, fMRI was chosen because it allows for unambiguous identification of subcortical sources of brain activity, which is crucial to the CMC and problematic for MEG analysis. The HCP includes functional neuroimages collected while participants performed seven psychological tasks. These tasks were taken or adapted from previously published influential neuroimaging studies and explicitly selected to cover the range of human cognition (Van Essen et al., 2013), therefore making it an ideal testbed for the *generality* criterion. Specifically, the tasks examine language processing and mathematical cognition (Binder et al., 2011), working memory, incentive processing and decision making (Delgado et al., 2000), emotion processing (Hariri et al., 2002), social cognition (Wheatley et al., 2007), and relational reasoning (Smith et al., 2007). The seven tasks were collected from six different paradigms (language processing and mathematical cognition were tested in the same paradigm).

To properly translate the CMC into a brain network architecture, its five components need to be identified with an equal number of spatially-localized but functionally homologous elements. Depending on the methods used, the number of *anatomically* identifiable areas in the human brain counts in the hundreds (Power et al., 2011; Yeo et al., 2011), and thus do not provide a reliable starting point. The number of *functionally* distinct circuits, however, is recognized as being at least one order of magnitude smaller, as different brain areas form interconnected networks (Cole et al., 2016; Power et al., 2011; Yeo et al., 2011). This study takes, as a reference point, the influential estimate given in Yeo et al. (2011), which counts seven distinct functional networks—a number that is comparable to the number of components in the CMC.

An initial identification can be made between CMC components and some of these networks. This initial identification was based on well established findings in the literature and is also consistent with the function-to-structure mappings that had been proposed in other neurocognitive architectures, such as the mappings suggested for ACT-R’s module-specific buffers (Anderson, 2007; Borst et al., 2015; Borst & Anderson, 2013) and the functional components employed in large-scale models of the brain (Eliasmith et al., 2012; O’Reilly et al., 2016). At this level, the working memory (WM) component can be identified with the fronto-parietal network comprising the dorsolateral prefrontal cortex (PFC) and posterior parietal cortex. The long-term memory (LTM) component corresponds with regions involved in the encoding of episodic memories, such as the hippocampus and the surrounding medial temporal lobe regions (Moscovitch et al., 2005; Squire, 2004), as well as with regions involved in memory retrieval, such as the medial frontal cortex and the precuneus; these regions are referred to as the default mode network (Raichle & Snyder, 2007). The action components can be identified with the sensorimotor network (Power et al., 2011); the procedural knowledge component with the basal ganglia (Yin & Knowlton, 2006); and the perception modules with the dorsal and ventral visual networks, as well as, depending on the task, the auditory networks (Figure 1B).

To properly characterize each individual component, a processing pipeline was designed to progressively identify a relevant corresponding region of interest (ROI) for each task and, within each task, for each of the ∼200 participants, thus accounting for individual differences in functional neuroanatomy (Figure 1C; See Materials & Methods).

### Alternative Architectures

To address our second criterion of *comparative superiority*, the CMC dynamic model was compared against other DCM models that implement alternative brain architectures.

Because the space of possible models is large, we concentrated on six examples that are representative of theoretical neural architectures previously suggested in the neuroscientific literature (Figure 2). These six alternatives can be divided into two families. In the “Hierarchical” family, brain connectivity implements hierarchical levels of processing that initiate with Perception and culminate with Action. In this family, the brain can be abstracted as a feedforward neural network model with large-scale gradients of abstraction (Huntenburg et al., 2018).

**Figure 2:**
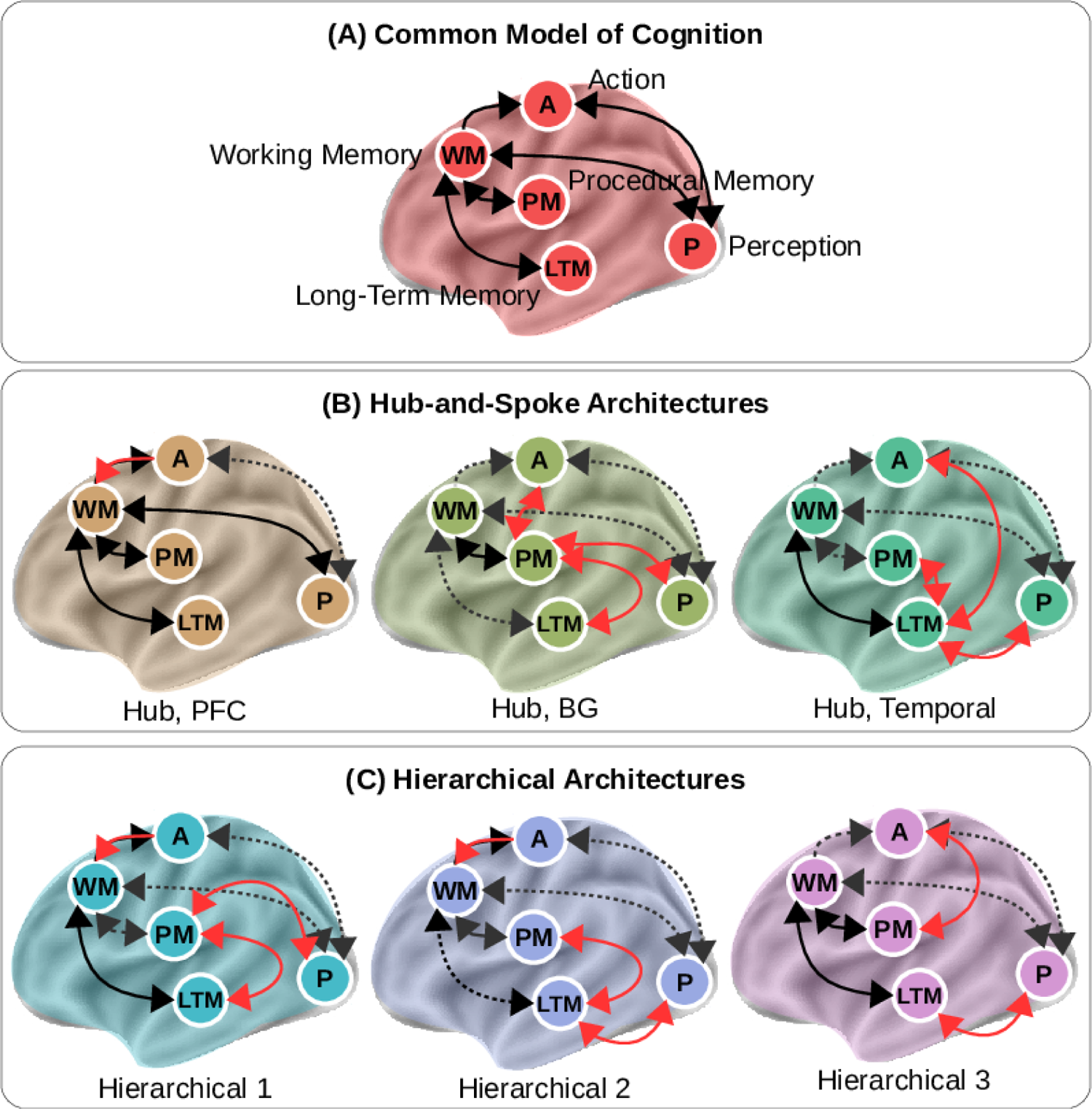
The seven architectures tested in this study. (A) The CMC; (B) The hub-and-spoke family of models, with the prefrontal (PFC), basal ganglia (BG), and temporal ROIs serving as the hub between the other modules; (C) The hierarchical family of models, representing three different configurations of working memory, procedural memory, and long term memory. In (B) and (C), pathways that are common to the CMC are shown in black; pathways that are present in the CMC but not included in the alternative models are shown as grey dashed arrows; and pathways that are present in the alternative models but not in the CMC are shown in red.

Within this hierarchical structure, each ROI represents a different level and projects both forward to the next level’s ROI and backward to the preceding level’s ROI. A degree of freedom in this architecture is the specific ordering of the regions within the hierarchy. Since Perception was always constrained to be the input and Action the output, the relative ordering of the remaining regions was manipulated. Furthermore, WM was considered as having a higher order in the hierarchy than LTM. With these constraints in place, the only remaining degree of freedom was the position of the Procedural region between Perception and Action, which gave rise to three possible hierarchical architectures (Figure 2B), in which the Procedural region falls between Perception and LTM (Hierarchical 1, as supported by models of the basal ganglia in perceptual categorization: Ashby, Ennis, & Spiering, 2007; Kotz, Schwartze, & Schmidt-Kassow, 2009; Seger, 2008), or between LTM and WM (Hierarchical 2, reflecting the role of basal ganglia in memory retrieval: Scimeca & Badre, 2012; Tricomi & Fiez, 2012), or between WM and Action (Hierarchical 3, as supported by models of basal ganglia in motor control Houk et al., 2007).

In the “Hub-and-Spoke” family (Figure 2C), a single ROI is singled out as the network’s “Hub” and receives bidirectional connections from all the other ROIs (the “Spokes”). With the exception of the Hub, no ROI is mutually connected to any other one. Three different Hub-and-Spoke architectures were created by selecting as the Hub one of the regions, with the exception of Perception and Action. In the first variant, the role of the Hub is played by the WM component. Because, in our mapping, the WM component corresponds to the lateral PFC, captures the view that the PFC functions as a flexible hub for control. This view is increasingly popular and well-supported by large-scale analysis of the human functional connectome (Cole et al., 2012, 2013). Interestingly, in terms of network architecture, this view is also the closest to the CMC, which, as noted above, is similarly based on a central WM hub, but also includes bidirectional Perception-Action connectivity. In the second variant, the role of the Hub is played by the Procedural Memory component, which reflects the centrality of procedural control in many production-system-based cognitive architectures (Anderson, 2007; Kieras & Meyer, 1997; Laird, 2012). Because, in our mapping, Procedural Memory is identified with the basal ganglia, this architecture also reflects the centrality of these nuclei in action selection and in coordinating cortical activity (Eliasmith et al., 2012; Hazy et al., 2007; Stocco et al., 2010). In the third variant, the role of the hub was played by LTM; this architecture reflects the convergence of cortical representations to form coherent semantic and episodic memories to interpret perception and guide action, and was the original namesake for “Hub-and-Spoke” (Chiou & Lambon Ralph, 2016; Rogers et al., 2004)

Thus defined, these six alternatives span all of the possible combinations within the Hierarchical and Hub-and-Spoke families, given the proposed constraints. Like the CMC, these architectures are representative of how the five components could be organized in a large-scale conceptual blueprint for the brain architecture; they simply make different choices as to which connections between components are more fundamental and better reflect the underlying neural organization. All of these architectures have been previously suggested in the literature as plausible plans to interpret the brain’s organization. In addition to representing plausible alternative architectures, these models differ minimally from the CMC and can be easily generated by replacing at most six connections from the CMC architecture (dashed lines and red lines, Figure 2B-C). Thus, any resulting differences in fit are unlikely to arise because of differences in network complexity.

### Modeling Network Dynamics

The link between the network of ROIs and their neural activity was provided through Dynamic Causal Modeling (DCM: Friston et al., 2003), a neuronal-mass mathematical modeling technique that approximates the time-course of brain activity in a set of brain regions as a dynamic system that responds to a series of external drives. Specifically, the time course of the underlying neural activity *y* of a set of regions is controlled by the bilinear state change equation:

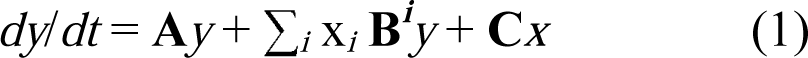

where *x* represents the event vectors (i.e., the equivalent of a design matrix in traditional GLM analysis), **A** defines intrinsic connectivity between ROIs, **C** defines the ROI-specific effects of task events, and **B** defines the modulatory effects that task conditions have on the connectivity between regions. For simplicity, the modulatory effects in **B** were set to zero, reducing the equation to the form **A***y* + **C***x*. A predicted time course of BOLD signal was then generated by applying a biologically-plausible model (the balloon model: Buxton et al., 1998; Friston et al., 2000) of neurovascular coupling to the simulated neural activity *y*.

Our preference for this technique was motivated by the existence of an integrated framework to design, fit, and evaluate models; by its ability to estimate the directional effects within a network (as opposed to traditional functional connectivity analysis); and by its underlying distinction between the modeling of network dynamics and the modeling of recorded imaging signals (as opposed to Granger causality), which makes it possible to apply the same neural models to different modalities (e.g., M/EEG data) in future work.

## Materials and Methods

The study presented herein consists of an extensive analysis of a large sample (N=200) of neuroimaging data from the Human Connectome Project, the largest existing repository of young adult neuroimaging data. The analysis was restricted to the task fMRI subset, thus excluding both the resting state fMRI data, the diffusion imaging data, and all of the M/EEG data. The task fMRI data consisted of two sessions of each of seven paradigms, designed to span different domains. All subject recruitment procedures and informed consent forms were approved by the Washington University in St. Louis’ Institutional Review Board. The present study met criteria for exemption at the University of Washington’s Institutional Review Board.

### Tasks fMRI Data

The HCP task-fMRI data encompasses seven different paradigms designed to capture a wide range of cognitive capabilities. Of these paradigms, six were included in our analysis; the Motor Mapping task was not included because it would have required the creation of multiple ROIs in the motor cortex, one for each effector (arm, leg, voice), thus making this model intrinsically different from the others. A full description of these tasks and the rationale for their selection can be found in the original HCP papers (Barch et al., 2013; Van Essen et al., 2013). This section provides a brief description of the paradigms, while Table 1 provides an overview.

**Table 1:**
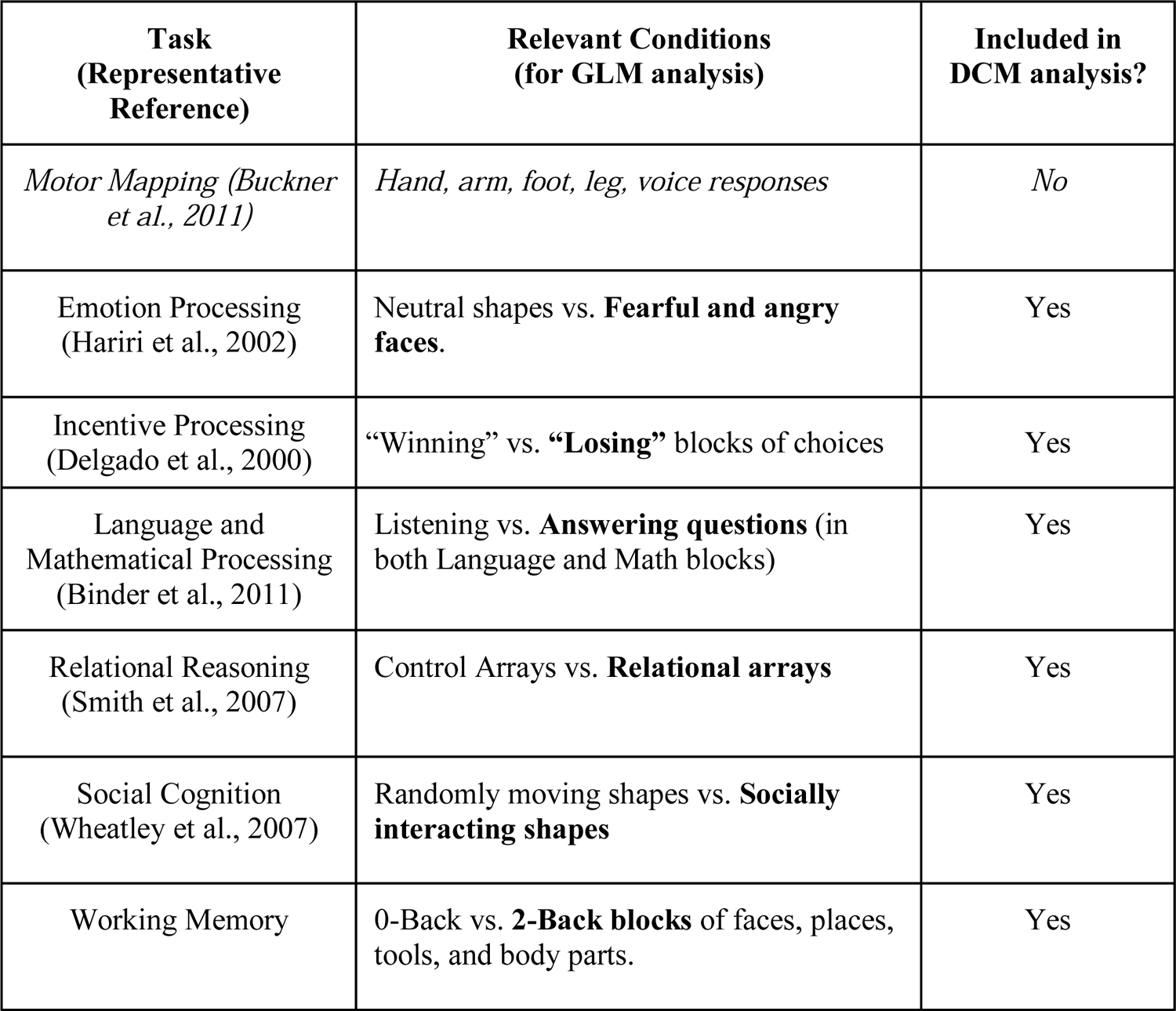
Overview of the seven task-fMRI paradigms used in the HCP dataset. Italics indicate tasks and conditions that were not included in our analysis; bold typeface marks experimental conditions that were selected as “Critical” (as opposed to “Baseline”) in the design of the experimental matrices (see below, “DCM-specific GLM analysis” section)

### Emotion Processing Task

Participants are presented with 12 blocks of six consecutive trials. During each trial, they are asked to decide which of two visual stimuli presented on the bottom of the screen match the stimulus at the top of the screen. In six of the blocks, all of the visual stimuli are emotional faces, with either angry or fearful expressions. In the remaining six blocks, all of the stimuli are neutral shapes. Each stimulus is presented for 2 s, with a 1 s inter-trial interval (ITI). Each block is preceded by a 3 s task cue (“shape” or “face”), so that each block is 21 s including the cue.

### Incentive Processing Task

The task consists of four blocks of eight consecutive decision-making trials. During each trial, participants are asked to guess whether the number underneath a “mystery card” (visually represented by the question mark symbol “?”) is larger or smaller than 5 by pressing one of two buttons on the response box within the allotted time. After each choice, the number is revealed; participants receive a monetary reward (+$1.00) for correctly guessed trials; a monetary loss (-$0.50) for incorrectly guessed trials; and receive no money if the number is exactly 5. Unbeknownst to participants, blocks are pre-designed to lead to either high rewards (6 reward trials, 2 neutral trials) or high losses (6 loss trials, 2 neutral trials), independent of their actual choices. Two blocks are designated as high-reward, and two as high-loss blocks. Each stimulus has a duration of up to 1.5 s, followed by a 1 s feedback, with a 1 s ITI, so that each block lasts 27 s.

### Language and Mathematical Processing Task

The task consists of 4 “story” blocks interleaved with 4 “math” blocks. The two types of blocks are matched for duration and adhere to the same internal structure in which a verbal stimulus is first presented auditorily, and a two-alternative question is subsequently presented. Participants need to respond to the question by pressing one of two buttons with the right hand. In the story blocks, the stimuli are brief, adapted Aesop stories (between 5 and 9 sentences), and the question concerns the story’s topic (e.g., “Was the story about *revenge* or *reciprocity*?”). In the math blocks, stimuli are addition or subtraction problems (e.g., “Fourteen plus twelve”) and the question provides two possible alternative answers (e.g., “*Twenty-nine* or *twenty-six*?”). The math task is adaptive to maintain a similar level of difficulty across the participants.

### Relational Processing Task

The task consists of six “Relational” blocks alternated with six “Control” blocks. In relational blocks, stimuli consist of two pairs of figures, one displayed horizontally at the top of the screen and one pair displayed at the bottom. Figures consist of one of six possible shapes filled with one of six possible textures, for a total of 36 possible figures.

Both pairs of figures differ along one dimension, either shape or texture; participants are asked to indicate through a button press if the top figures differ on the same dimension as the bottom figures (e.g., they both differ in shape). In the control blocks, the stimuli consist of one pair of figures displayed horizontally at the top of the screen, a third figure displayed centrally at the bottom of the screen, and a word displayed at the center of the screen. The central word specifies a stimulus dimension (either “shape” or “texture”) and participants are asked to indicate whether the bottom figure matches either of the two top figures along the dimension specified by the word. Both relational and control blocks have a total duration of 16 s, but they vary in the number of stimuli. Specifically, relational blocks contain four stimuli, presented for 3.5 s with a 500 ms ITI, while control blocks contain five stimuli presented for 2.8 s with a 400 ms ITI.

### Social Cognition Task

The task consists of 10 videoclips of moving shapes (circles, squares, and triangles). The clips were either obtained or modified from previously published studies (Castelli et al., 2000; Wheatley et al., 2007). In five of the clips, the shapes are moving randomly, while in the other five the shapes’ movement reflects a form of social interaction. After viewing each clip, participants press one of three buttons to indicate whether they believed the shapes were interacting, not interacting, or whether they were unsure. All clips have a fixed duration of 20 s with an ITI of 15 s.

### Working Memory Task

The task consists of eight 2-back blocks and eight 0-back blocks, with each block containing 10 trials. Each trial presents the picture of a single object, centered on the screen, and participants have to press one of two buttons to indicate whether the object is a target or not. In the 2-back blocks, a target is defined as the same object that had been seen two trials before, so that participants have to maintain and update a “moving window” of the past two objects to perform the task correctly. In the 0-back blocks, a target is defined as a specific object, presented at the very beginning of the block so that participants have to only maintain a single object in working memory throughout the block. The stimuli belong to one of four possible categories: faces, places, tools, and body parts. The category of the objects being used as stimuli changes from block to block, but is consistent within one block, so that there is an even number of face, place, tool, and body part blocks for each condition. Each block begins with a 2.5 s cue that informs the participant about the upcoming block type (2-back or 0-back). Each stimulus is presented for 2 s with a 500 ms ITI, for a total duration of 27.5 s per block.

### Data Processing and Analysis

#### Imaging Acquisition Parameters

As reported in Barch et al., (2013), functional neuroimages were acquired with a 32-channel head coil on a 3T Siemens Skyra with TR = 720 ms, TE = 33.1 ms, FA = 52°, FOV = 208 × 180 mm. Each image consisted of 72 2.0mm oblique slices with 0-mm gap in-between. Each slice had an in-plane resolution of 2.0 x 2.0 mm. Images were acquired with a multi-band acceleration factor of 8X.

#### Image Preprocessing

Images were acquired in the “minimally preprocessed” format (Van Essen et al., 2013), which includes unwarping to correct for magnetic field distortion, motion realignment, and normalization to the MNI template. The images were then smoothed with an isotropic 8.0 mm FWHM Gaussian kernel.

#### Canonical GLM Analysis

Canonical GLM analysis was conducted on the smoothed minimally preprocessed data using a mass-univariate approach, as implemented in the SPM12 software package (Penny et al., 2011). First-level (i.e., individual-level) models were created for each participant. The model regressors were obtained by convolving a design matrix with a hemodynamic response function; the design matrix replicated the analysis of Barch et al., (2013), and included regressors for the specific conditions of interest described in Table 1. Second-level (i.e., group-level) models were created using the brain-wise parametric images generated for each participant as input.

#### DCM-specific GLM Analysis

In parallel with the canonical GLM analysis, a second GLM analysis was carried out as part of the DCM analysis pipeline. The purpose of this analysis was two-fold. First, it defined the event matrix ***x*** that is used in the DCM equation (Eq. 1) to measure the parameter matrix **C**. Second, it provided a way to define the omnibus *F*-test that is used in the ROI definition (see below). Because these models are not used to perform data analysis, the experimental events and conditions are allowed to be collinear.

Like most cognitive neuroscience paradigms, each of our tasks includes at least two different conditions, under which stimuli must be processed in different ways. In all cases, the difference between conditions can be framed in terms of a more demanding, “critical” condition and an easier, “control” condition, with the more demanding events associated with greater mental elaboration of the stimuli. The critical condition of each task is emphasized in boldface in Table 1.

As is common in DCM analysis, these two task conditions were modeled in a layered, rather than orthogonal fashion. The difference is illustrated in Figure 3: While, in traditional GLM analysis, the two conditions are modeled as non-overlapping events in the design matrix, in the DCM-specific definition of the matrix all trials belong to the same “baseline” condition, which represents the basic processing of the stimulus across all trials. Stimuli from the critical condition form a subset of all stimuli presented in the baseline condition. The critical condition is therefore appended to the baseline condition in the design matrix to model the additional processes that are specifically related to it.

**Figure 3:**
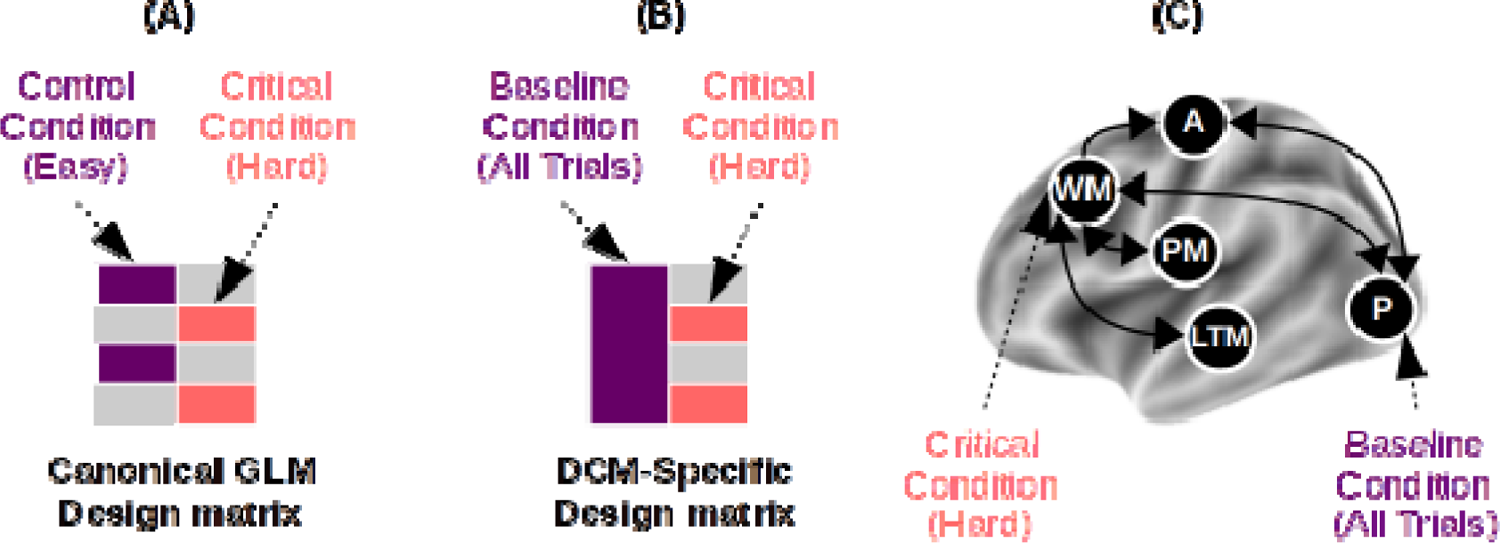
Difference between the design matrices used for canonical GLM (A) and for DCM analysis (B). In all of the network models, the Baseline condition drives neural activity in perceptual areas, while the Critical condition drives neural activity in the Working Memory component (C).

In DCM, each condition can affect one or more ROIs independently. In our analysis, the association between conditions and ROIs was kept constant across all tasks. Specifically, the baseline conditions selectively affected the perceptual ROI, while the critical condition selectively affected the WM ROI. This choice reflects the greater mental effort that is common to all critical conditions and is confirmed by the greater PFC activity found in all of the GLM analyses of the critical conditions (Barch et al., 2013).

### Regions-of-Interest Definition

To objectively define the Regions-of-Interest (ROIs) for each task and participant, a processing pipeline was set up. The starting point of the pipeline was an *a priori*, theoretical identification of each CMC component with large-scale neuroanatomical distinctions. As noted in the main text, this initial identification was based on well established findings in the literature as well as the function-to-structure mappings proposed in other large-scale neurocognitive architectures (Anderson, 2007; Borst et al., 2015; Borst & Anderson, 2013; Eliasmith et al., 2012; O’Reilly et al., 2016). Specifically, working memory (WM) was identified with the fronto-parietal network comprising the dorsolateral prefrontal cortex (PFC) and posterior parietal cortex; long-term memory (LTM) with regions in the middle, anterior, and superior temporal lobe; the procedural knowledge component with the basal ganglia; the action component with the premotor and primary motor cortex; and perception with sensory regions, including the primary and secondary sensory and auditory cortices, and the entire ventral visual pathway (Figure 1B).

Beginning with these macro-level associations, the pipeline progressively refined the exact ROI for each component through two consecutive approximations. Fig 1C provides a visual illustration of this procedure using the data from the relational reasoning task.

The first approximation was designed to account for group-level variability due to the different tasks and stimuli used in the four datasets. This was necessary because, for example, the different stimulus modalities determine which sensory area (e.g., auditory vs. visual areas) would be engaged and different task requirements would recruit different portions of the PFC. These differences were accounted for by conducting a separate group-level GLM analysis for each dataset, and identifying the coordinates of three points that have the highest statistical response within the anatomical boundaries of the visual areas (limited to the occipital lobe and the ventral portion of the temporal lobe), the dorso-lateral PFC, and the basal ganglia (limited to the striatum).

The second approximation was designed to account for individual-level variability in functional neuroanatomy. The group-level coordinates of each component, derived from the previous step, were then used as the starting point to search in 3D space for the closest active peak within the individual statistical parameter maps obtained from GLM models of each participant (see Figure 1C, right panel). For maximal sensitivity, the map was derived from an omnibus *F*-test that included all the experimental conditions. In practice, this *F*-test was designed to capture any voxel that responded to any experimental condition. The same *F*-contrast was also used to adjust (i.e., mean-correct) each ROI’s time series (Ashburner et al., 2016; Penny et al., 2011).

The individual coordinates, thus defined, were then visually inspected; when the coordinates were outside the predefined anatomical boundaries, they were manually re-adjusted. Across over 1,200 coordinates examined, only 2 required manual adjustment (∼ 0.2%). Figure 4 illustrates the distribution of the individual coordinates of each region for each task, overlaid over a corresponding group-level statistical map of task-related activity. Each individual coordinate is represented by a crossmark; the ∼200 crossmarks form a cloud that captures the spatial variability in the distribution of coordinates.

**Figure 4:**
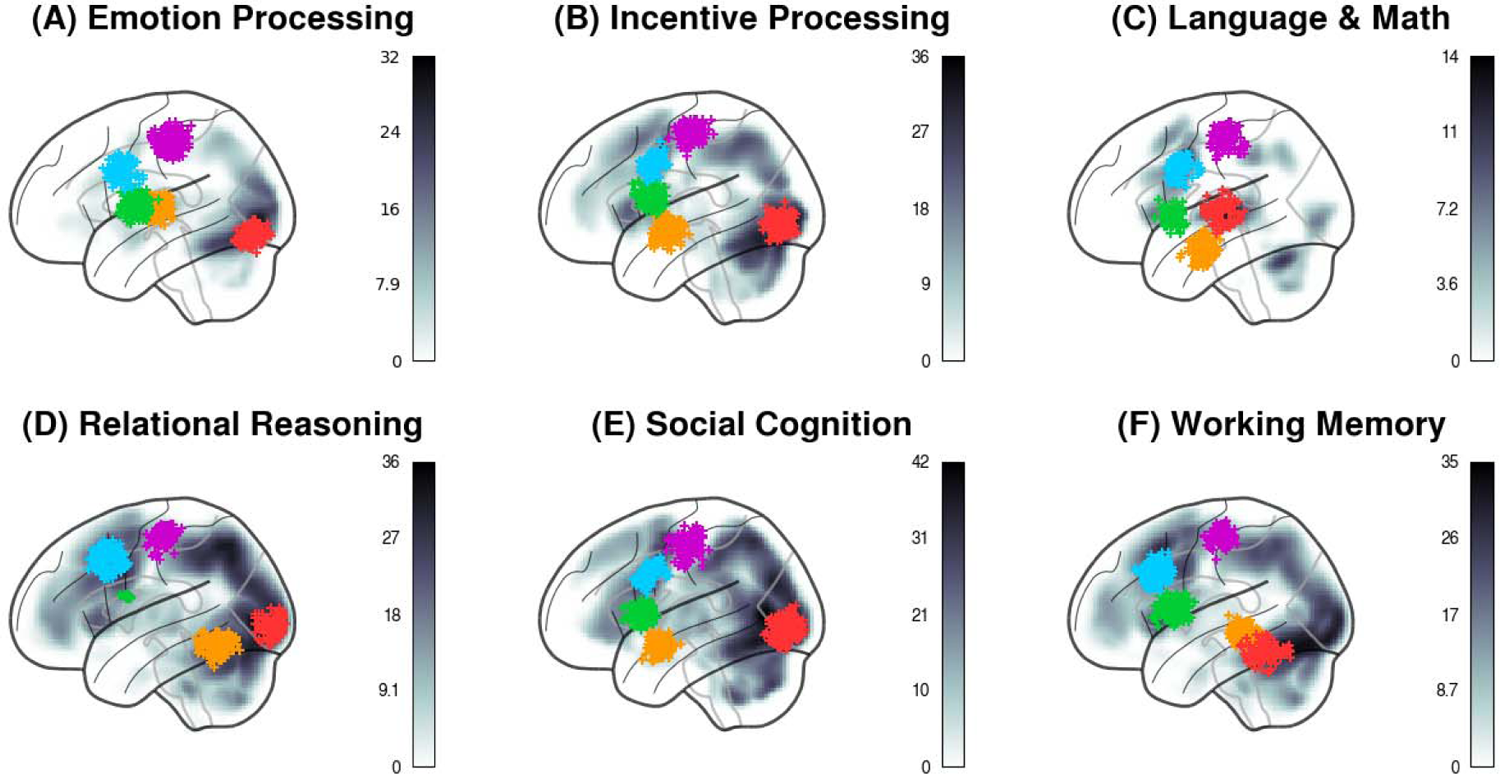
Lateral view of the distribution of the ROI centroids across individual participants and tasks. Each “+” marker represents the centroid of an ROI for one participant. Colors represent the components, following the conventions of *Fig 1A-C*. The background represents the statistical parametric map (in greyscale) of the corresponding group-level analysis used to identify the seed coordinates for each ROI (Step 2 in *Figure 1C*).

Finally, the individualized ROI coordinates were then used as the center of a spherical ROI. All voxels within the sphere whose response was significant at a minimal threshold of *p* < 0.50 were included as part of the ROI. For each ROI of every participant in every task, a representative time course of neural activity was extracted as the first principal component of the time series of all of the voxels within the sphere.

All the ROIs thus obtained were located in the left hemisphere: this simplifying approach was preferred to possible alternatives, such as including homologous regions in the right hemisphere (which would have required introducing additional assumptions about inter-hemispheric connectivity) or creating bi-lateral ROIs (which would have reduced the amount of variance captured in each ROI). Because all tasks show stronger activation in the left hemisphere than in the right, our results are still representative of brain activity in these domains.

### Model Fitting

Once the time-series for each ROI was extracted, different networks were created by connecting all of the individually-defined ROIs according to the specifications of each architecture (Figure 2). It should be noted that synaptic pathways exist that connect every pair of components; thus, this network model is designed to capture the fundamental layout of a brain architecture in terms of functionally necessary connections, rather than anatomical details.

The predicted time course of the BOLD response for each network model was then generated by using Equation 1 to simulate network activity as it unfolded over the course of the task. The predicted time course of BOLD signal was then generated by applying a biologically-plausible model (the balloon model: Buxton et al., 1998; Friston et al., 2000) of neurovascular coupling to the simulated neural activity y of each node in the network. The parameters of the full DCM, which include both the network connectivity parameters and the physiological parameters of the neurovascular coupling model, were estimated by applying the expectation-maximization procedure (Friston et al., 2003) to reduce the difference between the predicted and observed time course of the BOLD signal in each ROI.

## Results

Once the seven DCM models were separately fitted to the functional neuroimaging data, they were compared against each other using a Bayesian random-effects procedure (Stephan et al., 2009). Like many other model comparison procedures, this approach provides a way to balance the complexity of a model (as the number of free parameters) versus its capacity to fit the data. Compared to popular log-likelihood-based measures (e.g., Akaike’s information criterion: Akaike, 1974), this procedure is more robust in the face of outlier subjects, and thus better suited for studies that, like the present one, include a large number of participants and deal with considerable inter-individual variability (Stephan et al., 2010; Stephan et al., 2009).

Figure 5, inspired by Stephan et al., (2009), provides a graphical illustration of the procedure. Specifically, the probability *r_k_* that an architecture *k* would fit a random individual in a sample of participants is drawn from a Dirichlet distribution Dir(α, α,… α). This approach yields a posterior distribution of the probabilities *r_k_* for each model; the distributions of probabilities of architectures 1, 2,… *k* across *n* individuals are then drawn from multinomial distributions *m_k,n_* (see Figure 5A). Because of the properties of the Dirichlet distribution, the distributions *r_k_* will jointly sum up to one. Intuitively, these distributions can be thought of as the probability densities that a random participant will be best explained by a given architecture, and they must sum up to one because the space of architectures in a given comparison is finite and each participant must be best fit by one. Figure 5B-C illustrates a simple case with two hypothetical architectures, identified by the black and the grey lines, respectively. The figure depicts a case in which there is a high probability, centered at around *r_BLACK_* = 0.8, that any participant will be best fit by the first architecture (the black distribution). A second probability distribution (in grey), centered at around *r_GREY_* = 0.2, represents the probability that any participant will be fit by the alternative architecture. These two probability distributions can then be compared in terms of their relative *expected* and *exceedance* probabilities. The expected probability (red line in Figure 5B) is simply the mean of a distribution; again, the properties of the Dirichlet distribution guarantee that the sum of the means of all distributions is 1. In the example of Figure 5B, the mean for the black distribution is 0.74. The exceedance probability is the probability that the *r_k_* for a given architecture *k* is larger than the corresponding value of any competing models (Figure 5C). In the case of two possible architectures (*k* = 2), the exceedance probability can be easily calculated as the area of each distribution to the right of *r_k_* = 0.5. When more than two architectures are compared (*k* > 2), however, there are no straightforward closed-form solutions to derive the corresponding distributions’ exceedance probabilities. In this case, exceedance probabilities are calculated numerically by simulating 10,000 times the outcomes of sampling from the original distributions and computing the proportion of times each given architecture has the highest probability.

**Figure 5.**
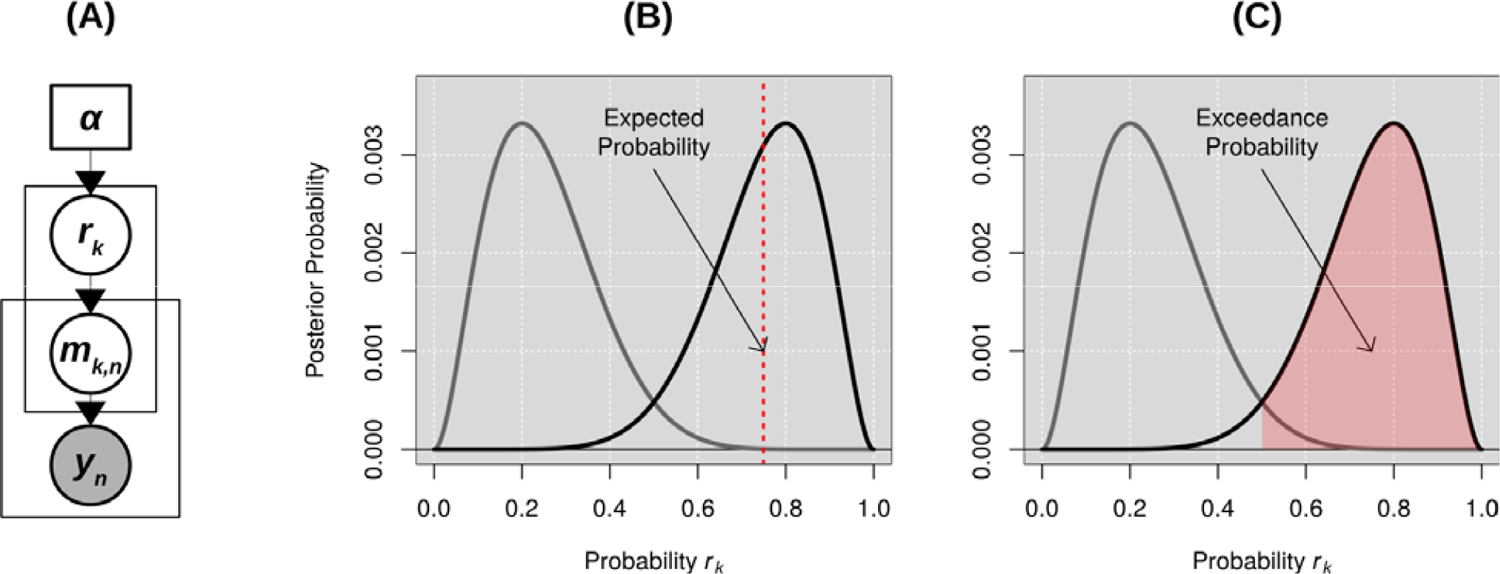
(A) Visual representation of the hierarchical Bayesian modeling procedure. (B) Visual representation of two architectures’ probability distributions r_k_, shown as the two thick grey and black curves. The red dashed line represents the expected probability of the winning architecture; (C) Visual representation of the winning architecture’s exceedance probability, that is, the proportion of a probability distribution that is greater than any other. In the case of two possible models (k = 2), the exceedance probability reduces to the area to the right of r_k_ = 0.5. Modified from Stephan et al., (2009).

The model posterior distributions are visualized for each task in Figure 6A-F. The expected probabilities are represented as the colored vertical lines, while the exceedance probabilities are summarized as colored bars in Figure 6H. Table 2 provides a detailed list of model comparison metrics, including the ones derived from the hierarchical Bayesian procedure used in this study α, expected, and exceedance probabilities) as well as the group-level log-likelihood of each model.

**Figure 6:**
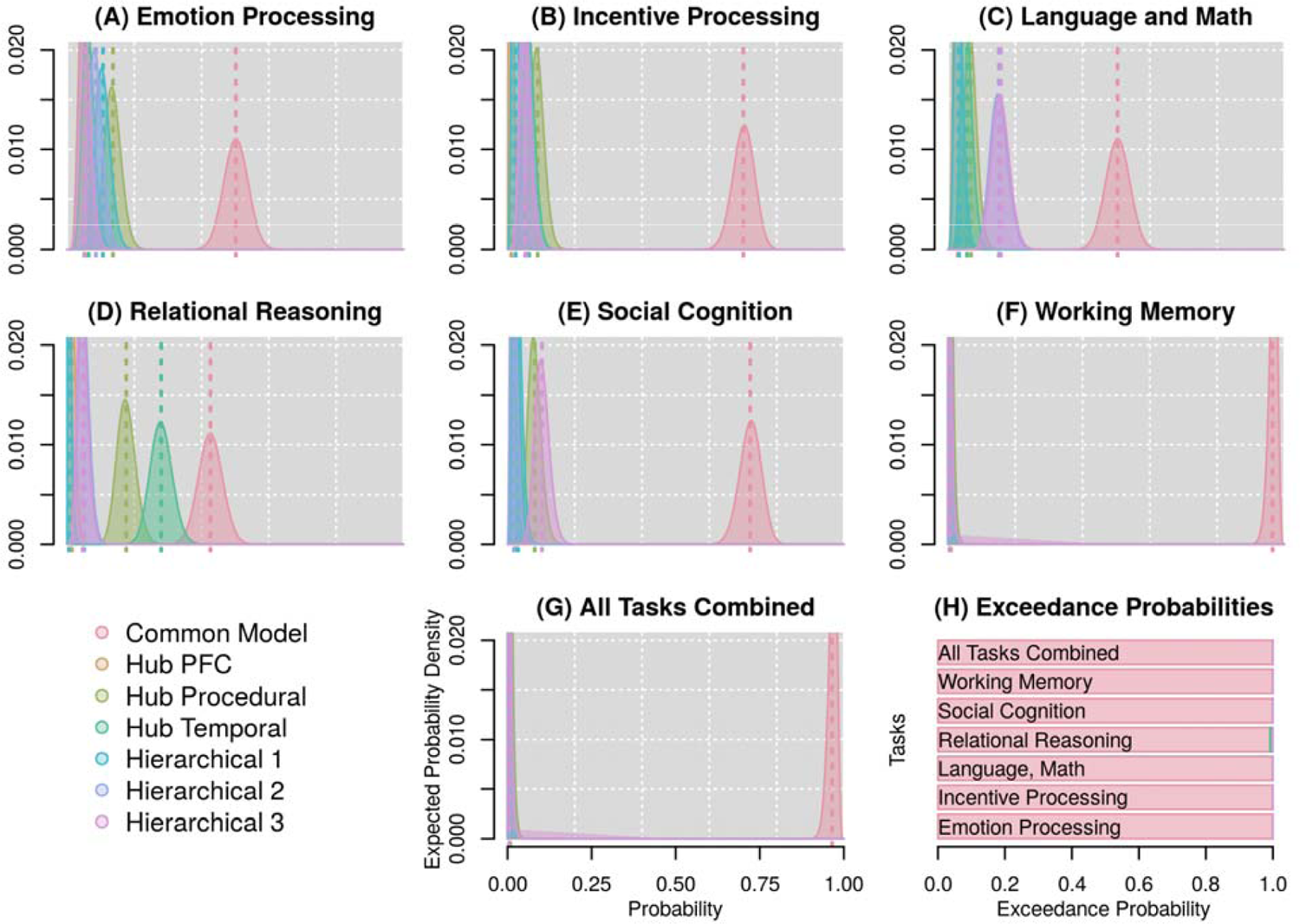
Results of the Bayesian model comparisons. In all plots, different colors represent different architectures. (A-G) Probability distributions that each of the seven architectures is true, given the data within each task and across all tasks combined. Vertical dotted lines represent the mean of each distribution, i.e. the expected probability of each model. (H) Corresponding exceedance probabilities, represented as stacked bars for each task.

**Table 2:**
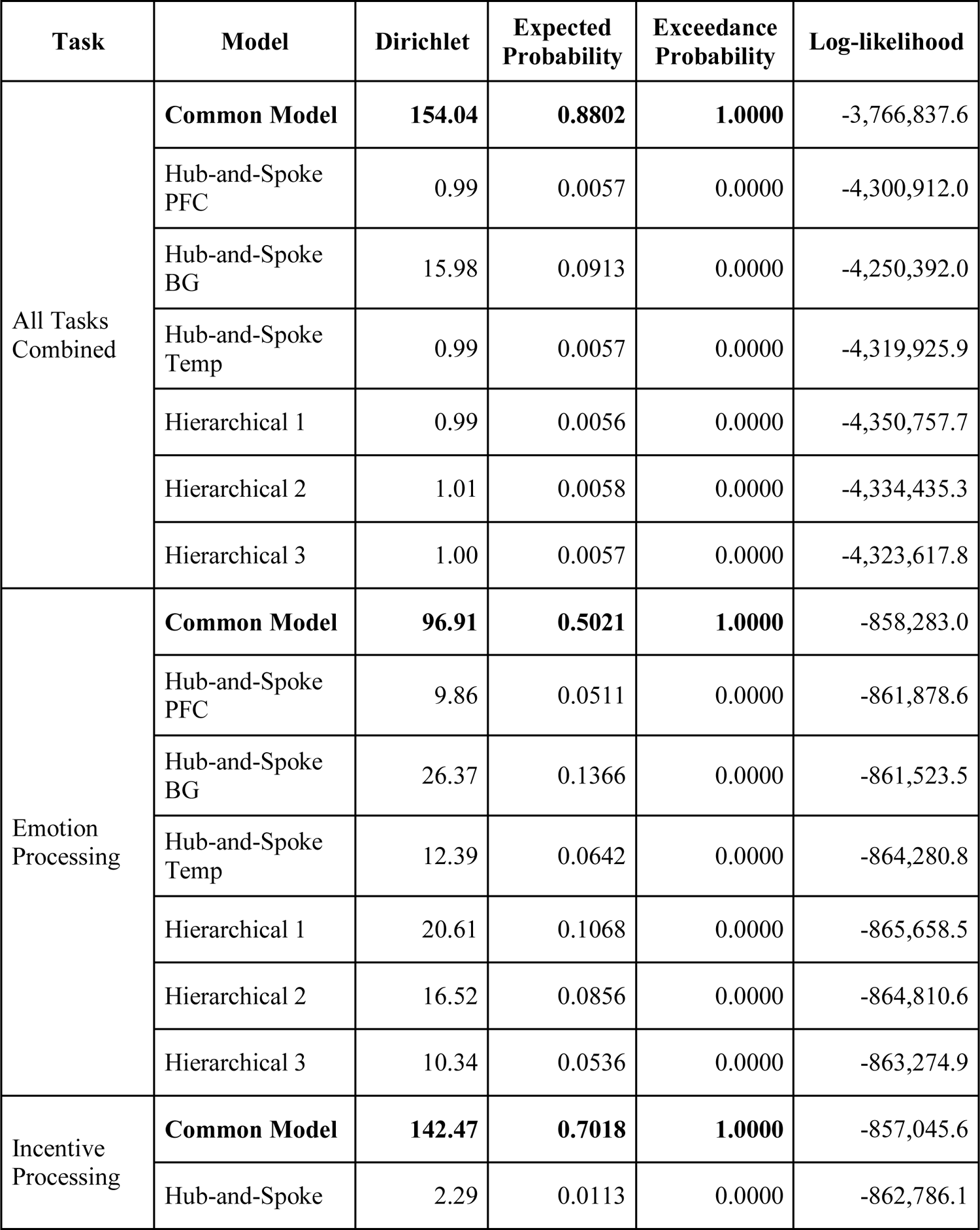
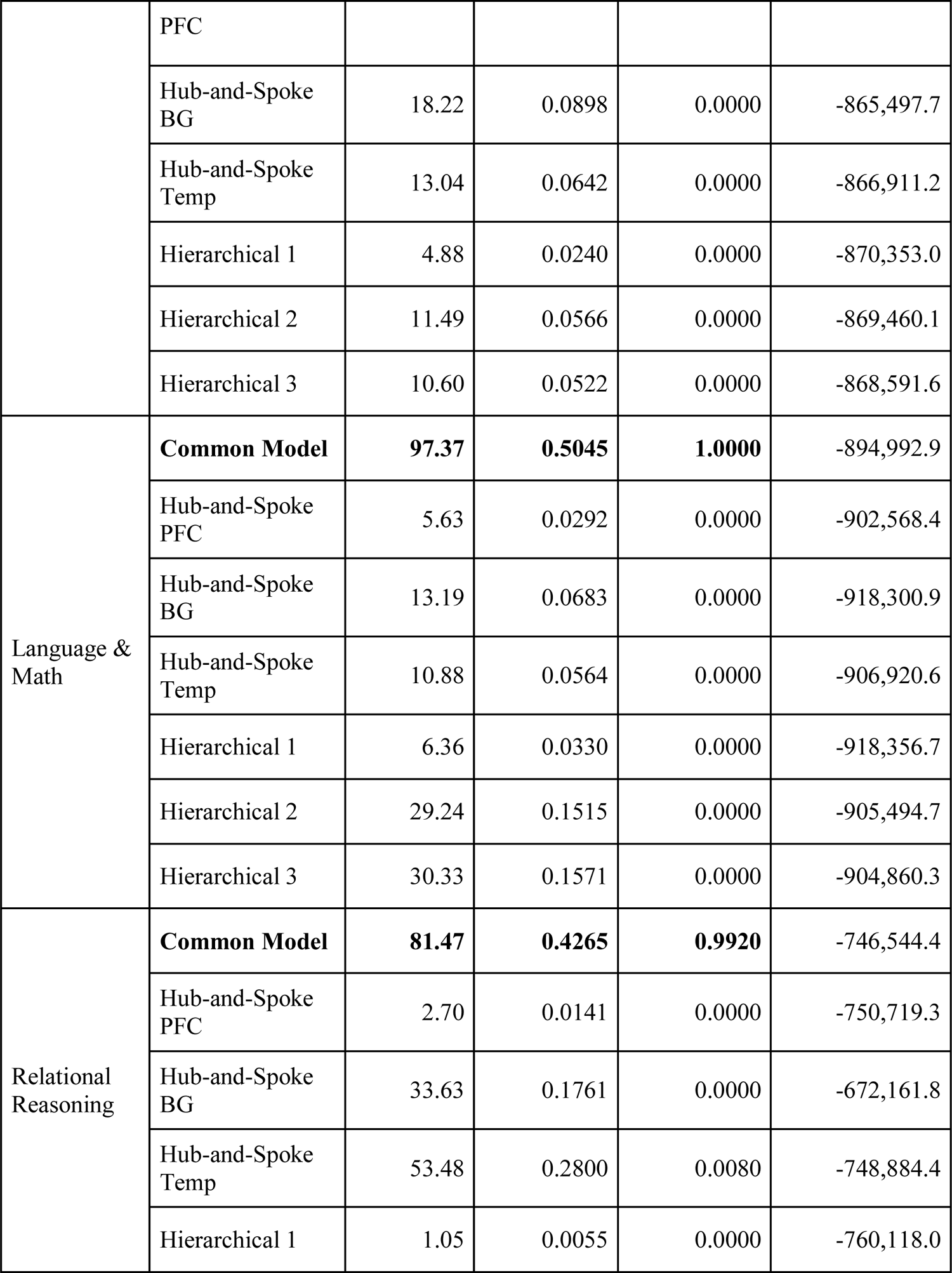
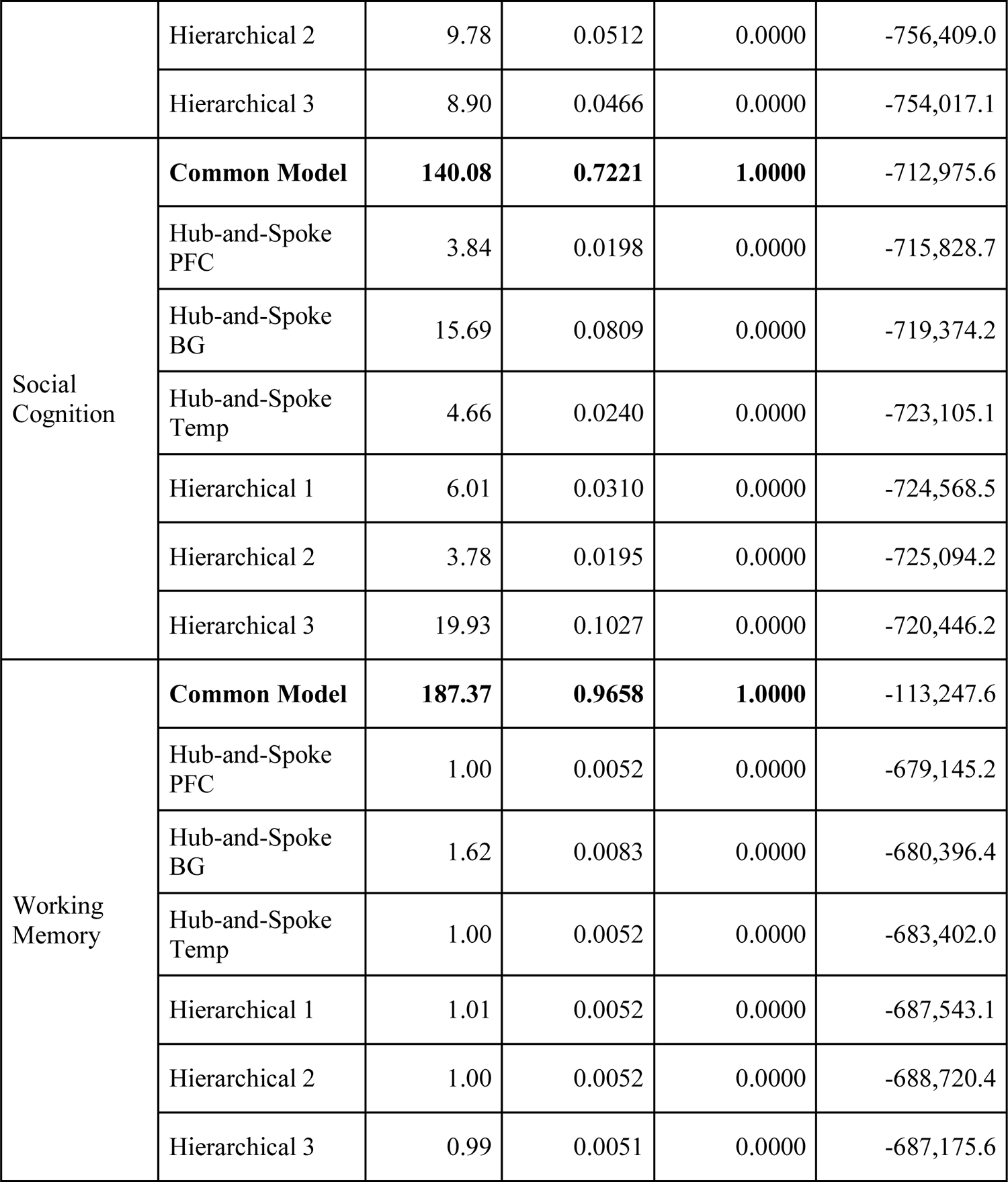
Results of Bayesian model comparison across tasks and models. For each task, the winning model is marked in bold.

Both types of metrics provide evidence in favor of the CMC. As shown in Figure 6A-F and Table 2, the CMC provides a better fit to the data than any alternative architecture, and its exceedance probabilities range from 0.99 to 1.0 (Figure 6H). Thus, the CMC uniquely satisfies both the generality and comparative superiority criteria. By contrast, all of the other architectures are consistently outperformed by the CMC in every domain (violating comparative superiority) and their relative rankings change from task to task (violating generality).

The only tasks in which another model comes close to the CMC in terms of fit were the Language and Mathematical cognition and the Relational Reasoning paradigms, in which the Hierarchical 2 model and the Hub Temporal model, respectively, reached an expected probability of 0.15 and 0.28 against the CMC (Figure 6H). Both paradigms stand out from the others for posing unusual demands in terms of switching between strategy or rules (Relational Reasoning) or between two entirely different tasks of comparable difficulty. This peculiarity raises a potential concern that the CMC’s superiority could be an artifact of modeling each task in isolation, and that in conditions where multiple tasks were modeled simultaneously, a different model could potentially provide a superior fit. To examine this possibility, a second analysis was carried out, which included only the 168 participants for whom data for all seven tasks was available. In this analysis, the data from each of the six paradigms performed by the same individual is modeled as a different run from a “meta-task” performed by that individual. When such an analysis was performed, the CMC maintained its superiority, all other models having a combined exceedance probability < 1.0 x 10^-10^ (Figure 6G-H, Table 2).

### The Role of Perception-Action Connectivity

One other possibility is that the superiority of the CMC originates from some peculiarity of its network connectivity that was missing in the other architectures. The one notable difference, in this sense, is the presence of a direct link between the Perception and Action ROIs, which are bilaterally connected in the CMC but unconnected in the six other rival architectures (Figure 2). To examine the role of a direct perception-action link in fitting the data, the six alternative architectures were augmented with bilateral Perception-Action connectivity and a new Bayesian model comparison was run. Notice that, after the addition of the Perception-Action links, the Hub Prefrontal architecture becomes virtually indistinguishable from the CMC,

differing only for the direction of one single connection (from the Working Memory to Action: Figure 2). Because, in a Bayesian model comparison, models compete against each other, two almost-identical models run the risk of evenly dividing the proportion of participants best explained, possibly leading to misleading low results. For this reason, we combined the two architectures into a single “family”, treating them as an identical model (Penny et al., 2010).

The results of these follow-up analyses are presented in Figure 7 and in Table 3 (for completeness, Table 3 reports the CMC and Hub-and-Spoke PFC entries separately). Overall, the addition of the direct Perception-Action connectivity improves the fit of the alternative models.

**Figure 7:**
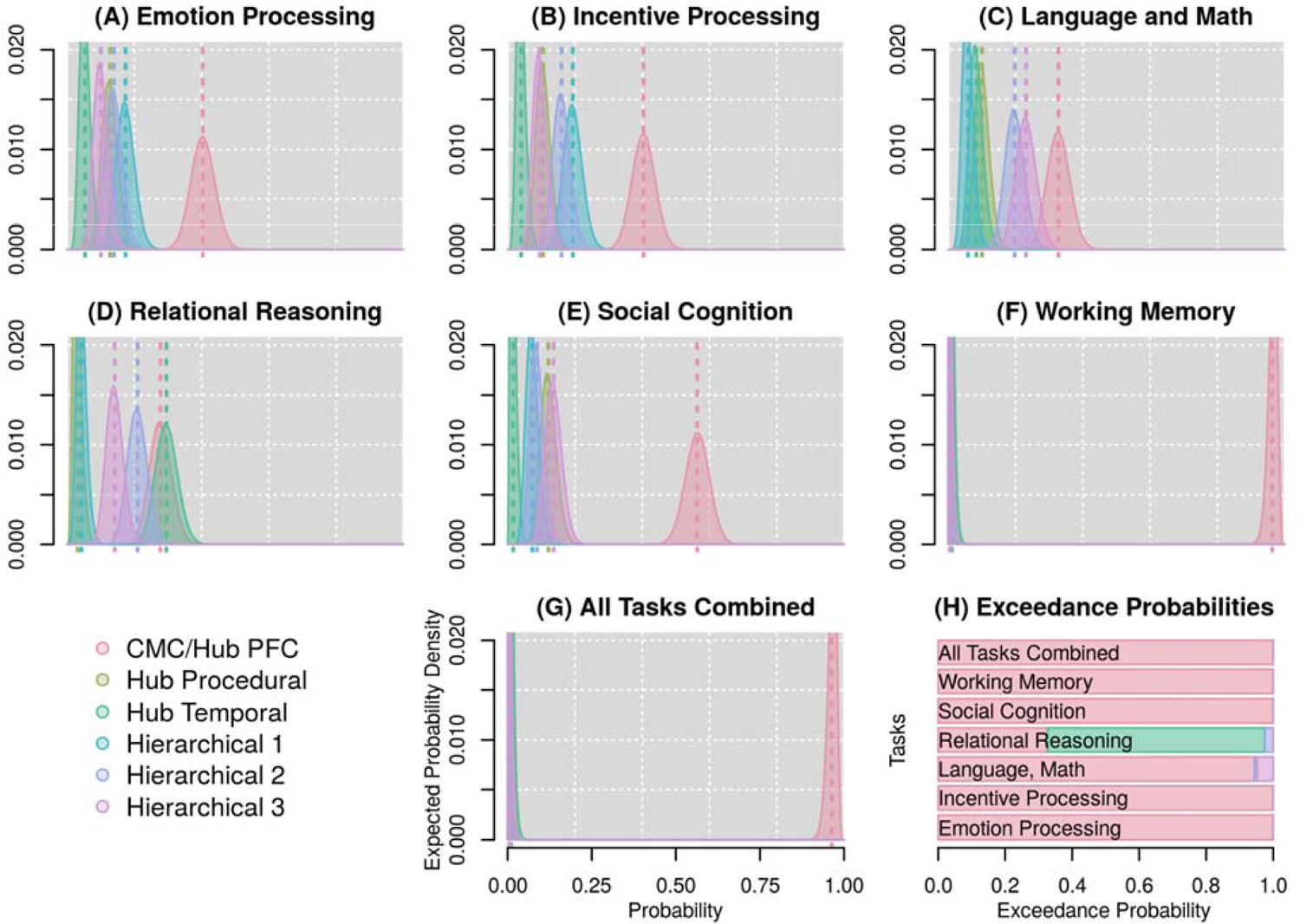
Follow-up Bayesian model comparisons, after the six alternative architectures have been augmented with bilateral Perception-Action connections. In all plots, different colors represent different architectures. (A-G) Probability distributions that each of the seven architectures is true, given the data within each task and across all tasks combined. Vertical dotted lines represent the mean of each distribution, i.e. the expected probability of each model. (H) Corresponding exceedance probabilities represented as stacked bars for each task.

**Table 3:**
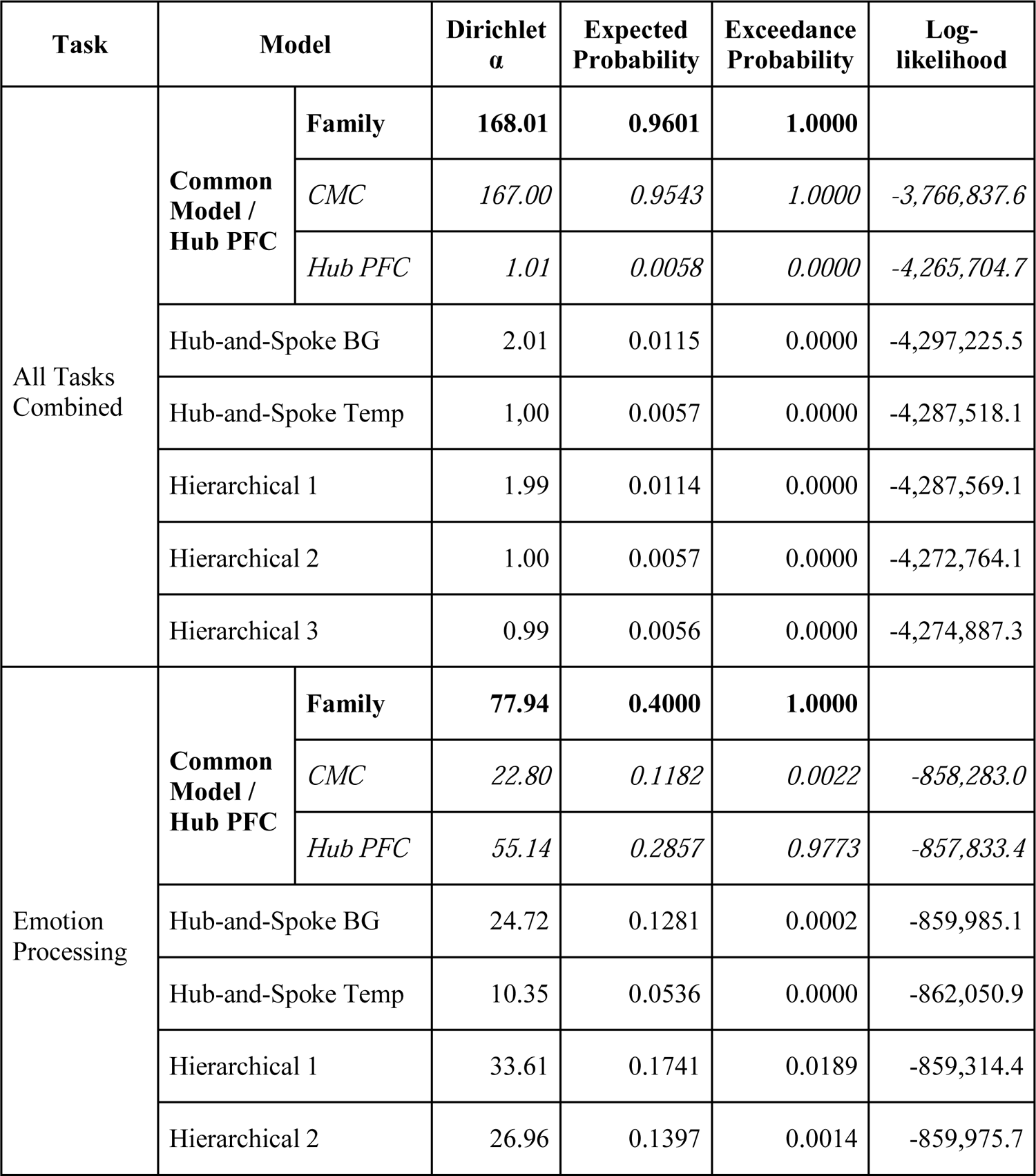
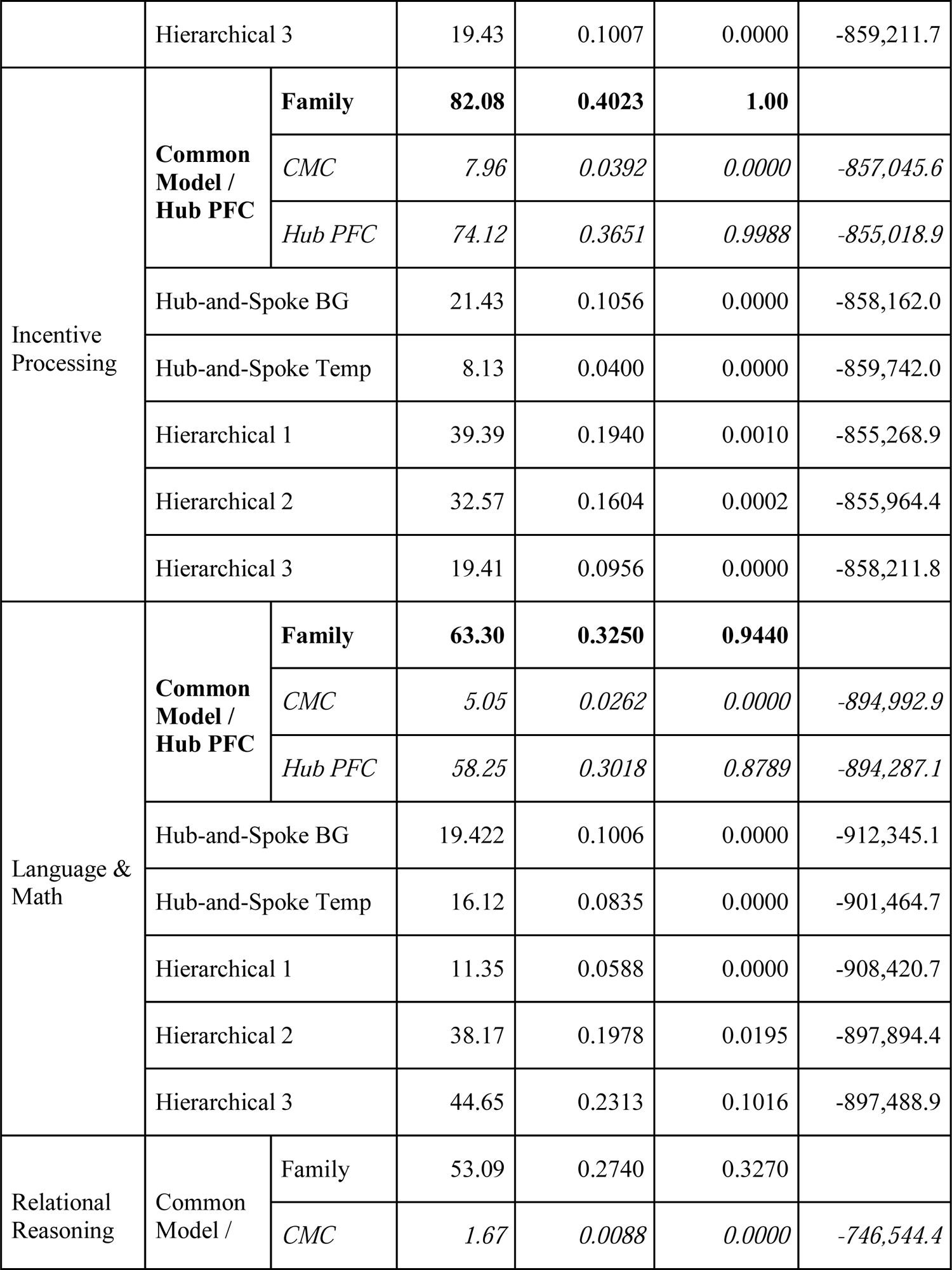
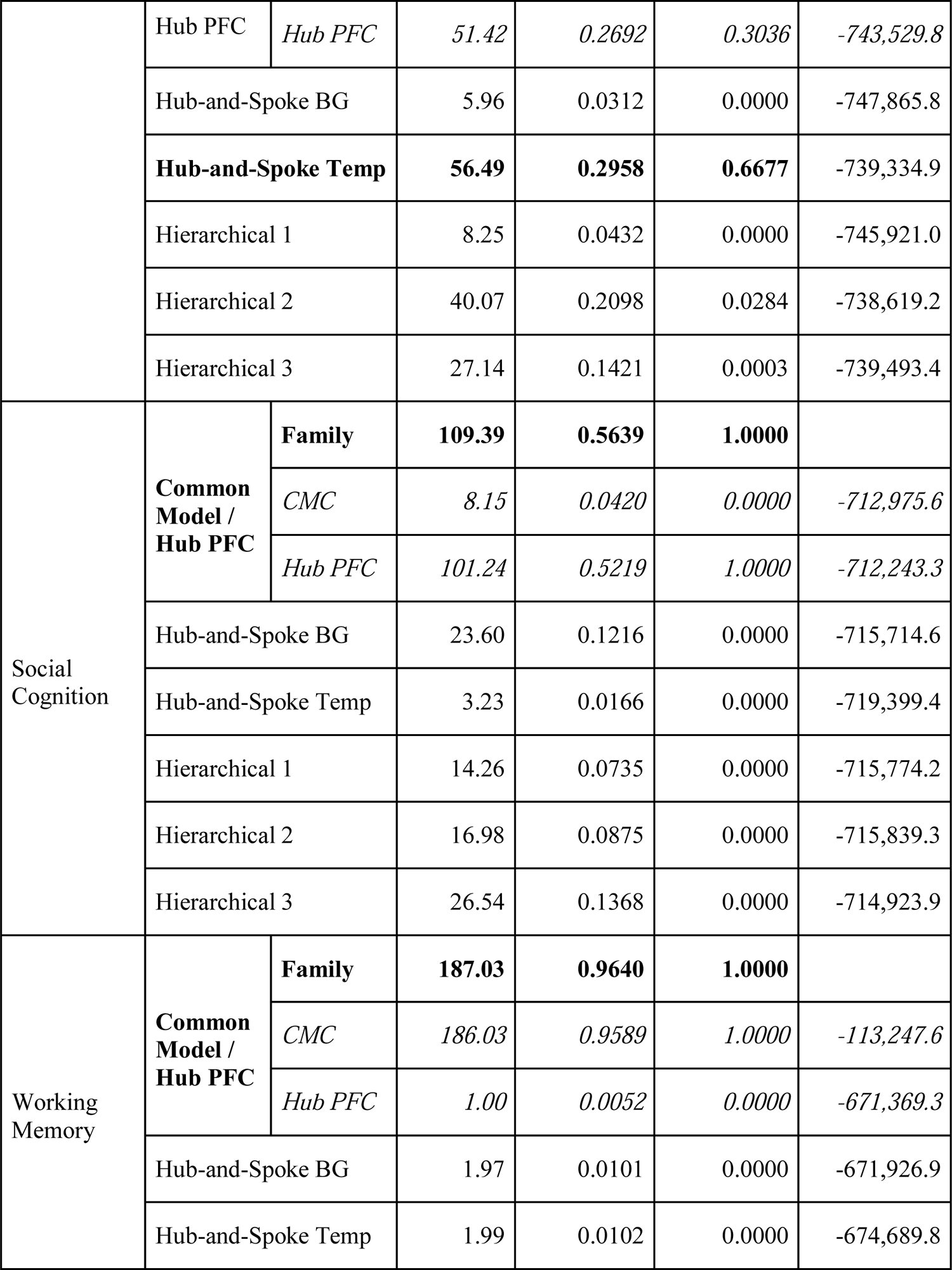
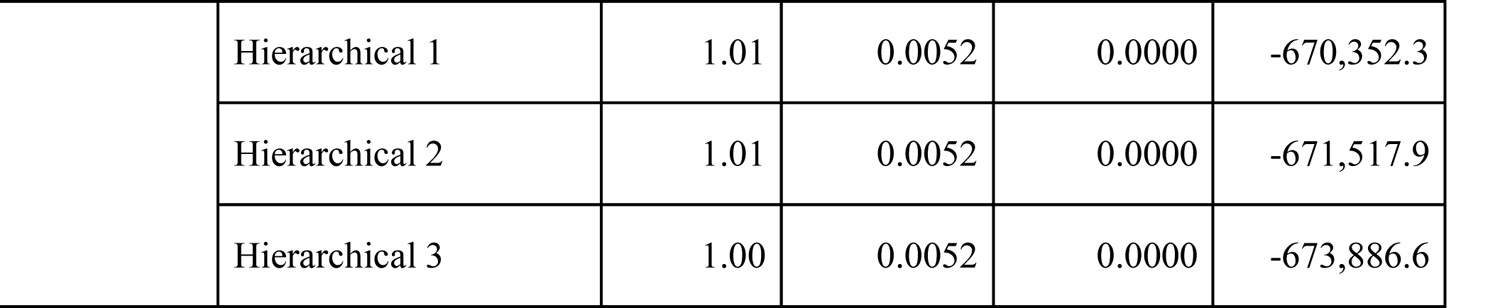
Results of Bayesian model comparison across tasks and models with added Perception-Action connection; the results of the Common Model of Cognition and the Hub-and-Spoke PFC architecture are presented separately. For each task, the winning model is marked in bold.

This improvement is shown in two ways. First, across all six tasks, the log-likelihoods of the alternative architectures have increased, on average, by 6,211; a fixed-effects ANOVA, using Task and Connectivity (with vs. without Perception-Action links) as factors, showed that the main effect of connectivity was significant [*F*(1, 60) = 6.411, *p* = 0.014], suggesting that their absolute fit to the data has grown reliably. Second, and more importantly, the expected probability of the alternative architectures has risen from *r* = 0.060 to *r* = 0.133, implying that their distributions have shifted rightwards. Once more, a Task-by-Connectivity fixed-effects ANOVA showed that this increase was significant [*F*(1, 60) = 12.42, *p* = 0.0008]. Because the expected probabilities need to sum to one, the growth of the alternative models must have occurred at the expense of the CMC, thus implying that the alternative models have become more competitive. This improvement notwithstanding, the results of the Bayesian model selection essentially replicates the previous analysis, showing that the CMC/Hub Prefrontal family provides a superior fit across tasks. Once more, the only exceptions are the Language and Math paradigm, where the Hierarchical 2 and 3 architectures also provide a reasonable fit (expected probabilities *r* = 0.20 and *r* = 0.23, respectively, against the CMC/Hub Prefrontal’s *r* = 0.32; Figure 7C and Table 3), and the Relational Reasoning task, where the Hub Temporal model provides, this time, a slightly better fit than the combined CMC/Hub Prefrontal architecture (expected probability of *r* = 0.30 vs *r* = 0.27; Figure 7D and Table 3). As in the previous case, a follow-up analysis was carried out by combining all tasks into a single paradigm and, thus, rule out the possibility that the combined CMC/Hub Prefrontal architecture would be underperforming under conditions that require integrating or switching between sources of information. Under this combined condition, the combined CMC/Hub Prefrontal family showed, once more, its superiority, dominating over all other models (expected probability *r* = 0.96; Figure 7G).

### Differences Between CMC and Augmented Hub Prefrontal

It has been noted several times that, once the Hub Prefrontal architecture is augmented with bi-directional Perception-Action connectivity, it becomes essentially indistinguishable from the CMC. This is because the difference between the two architectures reduces to the presence of one additional feedback connection (from the Action component to the WM component) in the Hub Prefrontal architecture; the absence of such connection, however, is *not* a central tenet of the CMC. Thus, while the differences between the remaining architectures are large, structural, and representative of different conceptual views of the brain’s organization, this difference is comparatively minor. It remains interesting, however, to consider whether it has functional implications.

In addition to reporting the data in which the two models are considered as a single family, Table 3 separately reports the relevant expected and exceedance probabilities and log-likelihoods for the CMC and augmented Hub Prefrontal. It is worth pointing out two relevant features in the data. The first is that, as expected, every time the joint CMC/Hub Prefrontal Family was selected as the best architecture, it was due to either one of its two resulting architectures. This is relevant because it could be argued that, by encompassing two different architectures (albeit very similar ones), this family was given an unfair advantage.

The second is that there is no consistent winner between the two architectures. Although the augmented Hub Prefrontal wins over the CMC in five out of six tasks (Emotion, Incentive Processing, Language + Math, Relational Reasoning, and Social Reasoning), the CMC vastly surpasses it in the Working Memory task. The degree by which the CMC outperforms in the working memory task is such that, when all tasks are combined together, the CMC again comes out as the best model. This is confirmed by an analysis of the two architectures’ log-likelihoods across task paradigms (Table 3). Unlike the exceedance and expected probabilities, which are constrained to sum up to one and thus depend on the other architectures in the comparison set, log-likelihoods are characteristic for each architecture. As the data in Table 3 indicates, the difference in log-likelihoods between the CMC and the augmented Hub Prefrontal is minimal across the five tasks in which the Hub architecture overperforms (mean difference = 1,382), but is massive in the Working Memory task (difference = 558,121), providing the decisive advantage when all tasks are combined.

### Analysis of CMC Connectivity

As noted earlier, although the competing architectures were chosen to represent current alternative views, we cannot entirely rule out the existence of alternative architectures that explain the data better than the CMC. It is possible, however, to decide whether all of the connections in the CMC are necessary, or whether a simpler model could potentially fit the data equally well. This is particularly important because, as noted in the previous section, the difference between the CMC, as outlined in the original paper (Laird, Lebiere, & Rosenbloom, 2017) and the modified Hub Prefrontal architecture (augmented with bilateral Perception-Action connections) boils down to a single, directed link.

To this end, a Bayesian parameter averaging procedure (Kasess et al., 2010) was conducted to generate the posterior distributions of the intrinsic connectivity parameter values (corresponding to matrix **A** in Eq. 1) across participants for each task. Figure 8 visually depicts the six task-specific connectivity matrices, indicating both the mean value (as the matrix cell color) and the associated posterior probability (as the overlaid number) for each CMC connection in each task. As the figure shows, the parameter values change significantly from task to task, implying that the CMC architecture is adaptively leveraged to meet the specific requirements of each paradigm. Nonetheless, virtually all parameters have a posterior probability *p* ≈ 1.0 of being different than zero (with just two out of 84 parameters having smaller posterior probabilities of *p* = 0.75 and *p* = 0.98), suggesting that all the components and their functional connections remain necessary across all domains.

**Figure 8.**
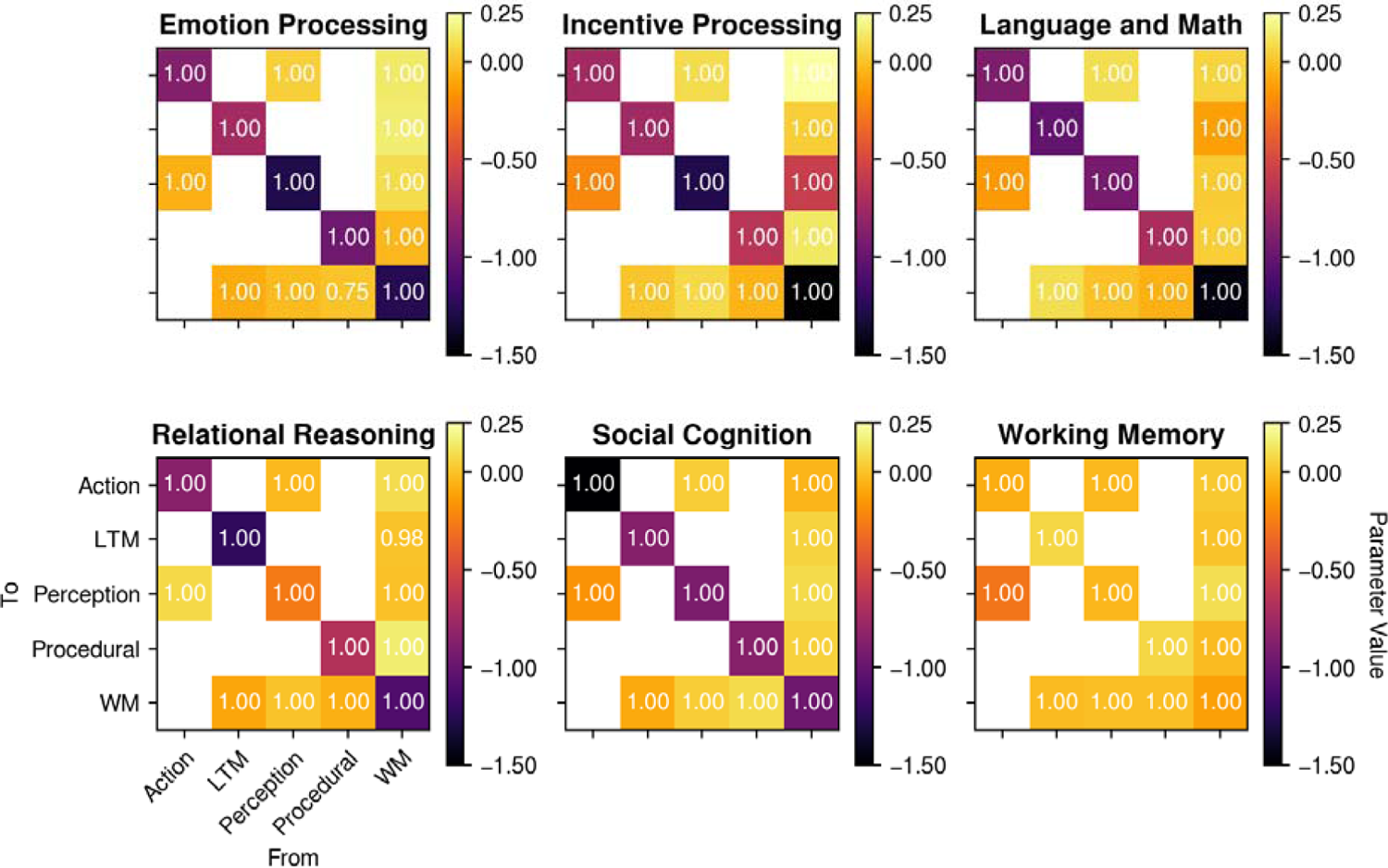
Estimated DCM intrinsic connectivity parameters for the CMC model. Each plot represents the intrinsic connectivity matrix (matrix **A** in Eq. 1); the cell color indicates the parameter value, and the white text indicates the posterior probability that the parameter value is significantly different from zero. White matrix cells indicate connections that are not present in the CMC (see Figure 1A).

## Discussion

In this study, a comparative analysis was performed on the relative ability of seven theoretical architectures to account for brain activity across seven different domains. These results provide overwhelming and converging evidence in favor of the Common Model of Cognition (CMC), a consensus architecture derived from the analysis of both human and artificial intelligent systems. Specifically, a CMC-inspired network model of the brain consistentlyoutperforms other architectures across all of the domains, thus jointly satisfying the two *a priori* criteria of *generality* and *superiority*. Even when the six alternative architectures were augmented with the CMC-specific bidirectional Perception-Action connectivity, either the CMC or a close variant remained dominant across both criteria. Thus, the CMC emerges as a viable high-level blueprint of the human brain’s architecture, potentially providing the missing unifying framework to relate brain structure and function for research and clinical purposes.

## Limitations

Although surprisingly robust and based on a large set of data, these results should be considered in light of four potential limitations. First, our conclusions are based on an analysis of task-related brain activity. Despite being established in the literature, the HCP tasks remain artificial and laboratory-tasks, and their ecological validity is, therefore, unknown. In contrast, many prominent studies have focused on task-free, resting-state paradigms (Cole et al., 2013, 2016; Fox et al., 2005; Power et al., 2011; Yeo et al., 2011). Thus, although the use of the task-related activity provides the most natural test for the generality criterion, the extent to which the CMC applies to resting-state fMRI remains to be explored.

Second, although this paper has argued that that the CMC components can be mapped to a small set of large-scale networks in the human brain (as identified in the works of Power et al., 2011; and Yeo et al., 2011), the identification between functional networks and components is not perfect; some components match with multiple networks and some networks have no clear mapping to the CMC. Thus, in the future, more theoretical work would need to be done to elucidate the connections between the large scale network organization of the human brain and the CMC.

Third, as noted above, our selection of alternative architectures was representative but not exhaustive. Although most of the distinct architectures that can be generated using just the five CMC components are likely to be unreasonable from a functional standpoint, some of them could, potentially, outperform the CMC. Furthermore, small variations remain possible within each architecture, and their functional role should be further explored.

Finally, it can be argued that our approach does not take full advantage of the possibilities of DCM, which makes it possible to accommodate non-linear, modulatory effects in the dynamic model. For example, the strategic role of procedural knowledge in the CMC (Assumption B3 of the original paper) is compatible with a “modulatory” view of the basal ganglia, which has also been argued for from a theoretical standpoint (Stocco et al., 2010) and empirically observed in at least one study using effective connectivity analysis (Prat et al., 2016). In this study, modulatory connections were deliberately not included, so to level the field and make the seven possible architectures more similar to each other in terms of overall complexity. Preliminary evidence, however, suggests that modulatory versions of the CMC might even outperform the standard version discussed herein (Steine-Hanson, Koh, & Stocco, A., 2019; Stocco et al., 2018).

## Consideration on the Nature of the CMC

These limitations notwithstanding, it is worth examining the possible implications of the CMC’s ability to account for neural data across cognitive domains. Our set of analyses suggests that the superiority of the CMC across cognitive domains stems from two key characteristics: the “Hub-and-Spoke” nature of its connectivity, with a central hub located in the prefrontal cortex, and the presence of a direct Perception-to-Action route. Neither of these two elements, in isolation, are responsible for the success of the CMC in explaining human neuroimaging data; the Hub-and-Spoke architecture alone, for instance, is outmatched not only by the CMC but by other architectures as well (Figure 6). By contrast, while the Perception-to-Action connectivity improves the relative competitiveness of all architectures, it does not change the superiority of the CMC (Figure 7). Thus, it is reasonable to conclude that both elements jointly concur in determining the success of the CMC. A possible functional explanation for this duality is that the two elements provide different and complementary routes for control. The hub-like nature of the PFC has been speculated upon for more than a century (Bianchi, 1922; Luria, 2012), and has been confirmed empirically by functional connectivity studies (Cole et al., 2012, 2013; Zanto & Gazzaley, 2013); computationally, it has been postulated that its existence creates a “global workspace” (Dehaene et al., 1998) in which information across different modalities can be integrated and shared, thus allowing effortful but advanced and flexible control over behavior— and, possibly, even consciousness (Dehaene & Naccache, 2001). Conversely, direct bi-directional links between Perception and Action systems in the human brain (Craighero et al., 1999; Fuster, 2004) are believed to support fast and automatic online monitoring and correction of motor behavior.

## Broader Implications

The results outlined here also have further implications. The first is that they highlight the fact that a greater degree of translation is possible, at the systems level, between the results of the cognitive sciences and those of the neuroscience and neuroimaging communities.

Specifically, they show that constraints developed at the levels of cognitive theories could be successfully translated into constraints at the level of the brain’s functional organization. Furthermore, it was shown that systems-level principles from the cognitive sciences are, in fact, compatible with an increasingly popular view about the hub-like nature of PFC in humans.

Our results also suggest that a different, top-down approach to the analysis of functional connectivity in the brain could be implemented. This approach could be extended in the future and integrated with the more common bottom-up approaches to the analysis of functional connectivity.

Finally, the fact that the CMC, which draws inspiration from high-level models of human cognition and *artificial* intelligent systems, accounts for the neural activity of the human brain, which is a low-level *biological* intelligent system, is also worthy of further consideration. In principle, solutions designed for artificial systems do not need to apply for biological systems, or vice versa. A mundane explanation is that this convergence simply reflects the fact that principles of brain organization have indirectly guided the development of cognitive architectures for human and machine intelligence (for instance, through the mediating influence of cognitive science). A more radical explanation is that the architectural space for general (or, at least, human-like) intelligence is inherently constrained and possibly independent of its physical realization, whether organic or artificial. In this sense, the CMC could be a model for *any* intelligent system, or at least for a large class of them, at different levels of organization. Both hypotheses are worth exploring in future research.

## Footnotes

Note that this meaning of “general intelligence”, as often used in artificial intelligence (Goertzel 2014) and cognitive science (Anderson & Lebiere, 2003), is different from what psychometricians intend as “general intelligence,” which is a hypothetical factor *g* explaining the person-level correlations between different tasks (Hunt, 2010). For a review of the relationship between these two meanings of “general intelligence,” as well as other definitions, see Legg & Hutter (2007).

